# Martini bead form factors for nucleic-acids and their application in the refinement of protein/nucleic-acid complexes against SAXS data

**DOI:** 10.1101/498147

**Authors:** Cristina Paissoni, Alexander Jussupow, Carlo Camilloni

## Abstract

Small-angle X-ray scattering (SAXS) use in combination with molecular dynamics simulation is hampered by its heavy computational cost. The calculation of SAXS from atomic structures can be speed up by using a coarse grain representation of the structure. Here following the work of Niebling, *et al.* (*J. Appl. Cryst., (2014), 47, 1190*) we derived the Martini beads form factors for nucleic acids and we implemented them, together with those previously determined for proteins, in the publicly available PLUMED library. We also implemented a hybrid multi-resolution strategy to perform SAXS restrained simulations at atomic resolution by calculating on-the-fly the virtual position of the Martini beads and using them for the calculation of SAXS. The accuracy and efficiency of the method is demonstrated by refining the structure of two protein/nucleic acid complexes. Instrumental for this result is the use of metainference that allows considering and alleviating the approximations at play in our SAXS calculation.

## 1. Introduction

Small-angle X-ray scattering (SAXS) is a powerful structural technique to study biomolecules in a solution environment. Even if it does not reach atomic resolution, SAXS can complement and be integrated with other structural techniques providing information on the size, shape, global dynamics and intermolecular interactions of a system (Tuukkanen *et al.*, 2017; Koch *et al.*, 2003). Furthermore, time-resolved SAXS measures can be employed to study conformational changes over multiple time scales (Levantino *et al.*, 2015).

From a computational perspective calculating SAXS given a structure of *N* atoms is an O(*N*^2^) problem. In comparison, the calculation of NMR observables like chemical shifts, ^3^J-couplings or residual dipolar couplings require only few atoms (Schwieters *et al.*, 2003). This poses some limitation on the use of SAXS as a restraint or as a scoring function for large systems and large numbers of conformers.

Multiple strategies have been adopted to calculate SAXS efficiently, for example *CRYSOL* adopted a spherical harmonics expansion (Svergun *et al.*, 1995), while other approaches include hierarchical algorithms (Berlin *et al.*, 2014) or the particle mesh Ewald summation (Marchi, 2016). Alternatively, a possible strategy consists in coarse graining the structure representation using *M* beads with *M*<*N*, each comprising a variable number of atoms (Yang *et al.*, 2009; Ravikumar *et al.*, 2013; Stovgaard *et al.*, 2010; Zheng & Tekpinar, 2011; Niebling *et al.*, 2014). Recently, Niebling *et al.* (2014) derived the Martini beads form factors for proteins making use of the single bead approximation (SBA) (Yang *et al.*, 2009) and showed how this approach can be almost 50 times faster than the standard SAXS calculation, while retaining good accuracy for *q* values up to 0.5 Å^−1^.

Here, we first build on the work of Niebling, *et al* by deriving the Martini beads (Uusitalo *et al.*, 2015, 2017; Marrink & Tieleman, 2013) form factors for nucleic acids (DNA and RNA); then we implement a hybrid multi-resolution strategy where the position of Martini beads is calculated on-the-fly in a full atomistic molecular dynamics (MD) simulation and employed in combination with the abovementioned form factors to calculate SAXS. We demonstrate the accuracy and efficiency of this approach by refining the structure of a protein-DNA and a protein-RNA complex using the measured SAXS as restraints. Importantly, in our strategy, the strength of the restraint is determined by metainference, a Bayesian inference approach that allows considering multiple sources of errors, in such a way that the approximations at play are considered (Bonomi *et al.*, 2016). The presented approach, including the form factors for Martini beads of proteins and nucleic acids, is implemented in the PLUMED-ISDB module (Bonomi & Camilloni, 2017) of the PLUMED library (Tribello *et al.*, 2014), making it readily available to all the codes compatible with PLUMED as well as a standalone analysis tool.

## 2. Theory and Methods

### 2.1. Computing scattering intensities of biomolecules in solution

Given a molecule of *N* atoms the total scattering amplitude at vector ***q*** is described by:

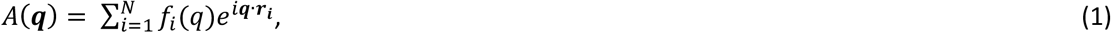

where ***r**_i_* and *f_i_* denoted the position and atomic scattering factor of atom *i*, respectively. If molecules are randomly oriented, it is possible to use of the Debye equation to compute the scattered intensity:

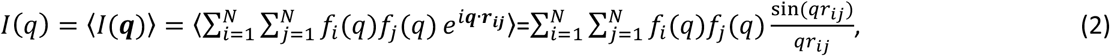

where *q* = |***q***| = 4*π* sin *θ/λ*, being 2*θ* the scattering angle and *λ* the X-ray wavelength, ***r**_ij_* is the vector distance from particle *i* to *j* and 〈…〉 indicates the spherical average over all the orientations. Here, the atomic scattering factor *f_i_*(*q*) can be computed using the Cromer-Mann analytic function:

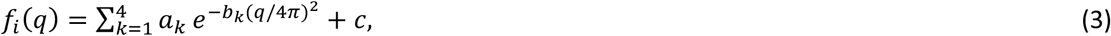

with the *a_k_, b_k_* and *c* parameters available in the *International Tables for Crystallography* (Cromer & Waber, 1965; Brown *et al.*, 2006).

While equation (2), in combination with these form factors, is effective in computing scattering intensities of biomolecules in vacuum, additional effects must be considered for a realistic representation of scattering in solution: 1) the electron density of the solvent displaced by the molecule; 2) the excess of electron density in the hydration shell; and 3) the conformational averaging of the molecules.

In SAXS measures, the background buffer scattering is subtracted from the sample scattering to remove unwanted solvent signal. To take into account the displaced solvent effect, an approach commonly adopted consists in using reduced atomic scattering factors, according to Fraser *et al.* (1978):

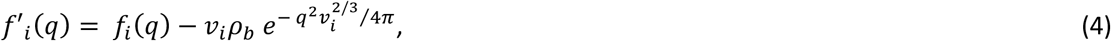

where *v_i_* is the tabulated displaced solvent volume of atom *i* and *ρ_b_* is the electron density of bulk water (e.g. 0.334 *e* Å^−3^). These modified form factors can be used in place of *f_i_*(*q*) in the Debye equation to include the effect of the displaced solvent.

Further, it should be considered that the density of the solvent layer around molecules can be different from the density of bulk solvent due to solute-solvent interactions, thus resulting in additional scattering terms. The inclusion of this effect in theoretical calculations of scattering intensities can be performed either modelling the hydration shell implicitly, as in *CRYSOL* (Svergun *et al.*, 1995) and FoXS (Schneidman-Duhovny *et al.*, 2010) among the others, or via more computationally expensive explicit-solvent approaches (Chen & Hub, 2015; Knight & Hub, 2015; Chen & Hub, 2014; Köfinger & Hummer, 2013; Park *et al.*, 2009). Most of these techniques requires to adjust one fitting parameter against experimental data, in order to tune the level of contrast in the hydration shell. Interestingly, it has been shown (Björling *et al.*, 2015; Niebling *et al.*, 2014) that if data are recorded as differences between two states, which is the typical case in time-resolved scattering experiments, the contribution of the solvation layer could be neglected.

Finally, conformational averaging can be included by averaging over multiple configurations of the system generated for example by MD simulations (Yang *et al.*, 2009).

### 2.2. Coarse-grain form factors

The Debye equation (2) requires the evaluation of pairwise distances between all the atoms in a biomolecule. This is an *O*(*N*^2^) problem, where *N* is the number of atoms, that becomes more and more computationally expensive as the dimension of the system increases. This is particularly serious when multiple evaluations of the scattering profile are required, as in the case of MD simulations driven by SAXS data; in iterative refinement and modelling; or when several trial structures must be tested. Several approaches circumvent this problem adopting a coarse-grain representation of the biomolecule to reduce the cost of the Debye summation (Yang *et al.*, 2009; Ravikumar *et al.*, 2013; Stovgaard *et al.*, 2010; Zheng & Tekpinar, 2011; Niebling *et al.*, 2014). According to this strategy, which is well justified by the low resolution of SAXS data, the molecule of interest is represented as a collection of beads, each comprising a variable number of atoms. The dimension of the beads can be tuned to find a proper balance between accuracy and computational efficiency (Niebling *et al.*, 2014). Beads can also be placed in different positions: examples include the atoms centre of mass, the centre of electron density distribution or, in the case of protein residues, the C*α* atom (Tong *et al.*, 2016). Given *M* beads and their associated scattering factors *F*(*q*), the Debye equation becomes:

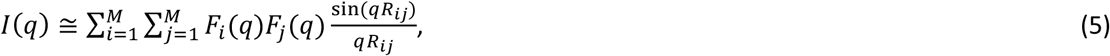

where the indices *i* and *j* run over the beads and *R_ij_* is their relative distance. Computing accurate coarse-grain scattering factors *F*(*q*) is a non-trivial task and diverse strategies, with different degrees of accuracy, can be employed. A review and a comparison of some of these possibilities, accompanied with a description of the approximations used in each case, is given in Niebling *et al.* (2014). Among these, the SBA method proposed by Yang *et al.* (2009), emerged as a reliable and fast method to calculate effective form factors. Herein *F_i_*(*q*) is calculated to reproduce the scattering intensity of the isolated bead *i* according to:

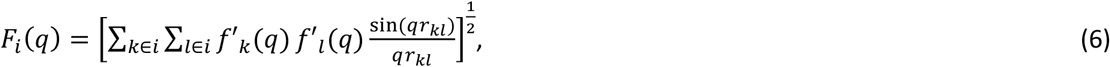

where the atomic scattering factors are the ones corrected for the excluded volume. The use of the reduced form factors in (6) could cause the violation of the condition:

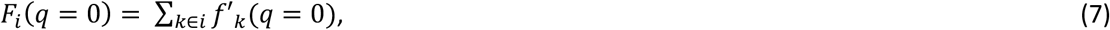

which guarantees to compute correct scattering intensities at *q*=0. This is due to the fact that equation (6) always produces positive coarse-grain form factors, while the sum in equation (7) can be negative (e.g. in the case of small beads containing several hydrogen atoms, whose reduced atomic scattering factor at *q*=0 corresponds to −0.72 electron units). To overcome this problem Niebling *et al.* (2014) proposed to correct the form factors *F_i_*(*q*) not satisfying equation (7) by fitting a sixth-order polynomial to data with *q* larger than the high-*q* inflection point and imposing a constraint at *q*=0. The resulting curve is then used as the corrected coarse-grain form factor.

### 2.3. Coarse-grain nucleic acids representation with Martini

Here, we applied the SBA approach to compute coarse-grained form factors for DNA and RNA, using as mapping scheme the Martini force field (Uusitalo *et al.*, 2015, 2017; Marrink & Tieleman, 2013) and following the work done by Niebling *et al.* (2014) for proteins. In the Martini force field, each nucleotide is represented with six or seven beads. The backbone is mapped with three beads (one including the phosphate group and two comprising the atoms of the sugar ring), while the nitrogen bases are modelled with three (cytosine and thymine) or four beads (adenine and guanine).

We computed coarse-grain SBA form factors for each Martini bead in the eight nucleotides, averaging over a large number of crystal structures. The correction described in the previous section was applied for those beads not satisfying equation (7) (i.e. the backbone beads BB3 of DNA nucleotides, see Table S1). Furthermore, we added two terminal beads for each nucleotide. The terminal beads are denoted as TE5 and TE3 (for the 5’ and 3’ termini) and are obtained including the terminal hydroxyl groups into the backbone beads BB2 and BB3, respectively. In the calculation of scattering intensities, the position of each bead was placed in the centre of mass of the nonhydrogen atoms belonging to the bead, according to the Martini model. The only exception is represented by the positioning of the termini, for which the terminal oxygen atom was not considered to be coherent with the Martini representation of the nucleotide. This implies that Martini structures can be directly used as input for the evaluation of scattering intensities, facilitating the use of SAXS data as restraints in Martini MD simulations.

### 2.4. DNA and RNA data sets

In order to compute accurate coarse-grain form factors considering the internal details of each bead, the summation in equation (6) is expected to be averaged over a number of different conformations. To achieve this, a set of non-redundant molecular structures from the Protein Data Bank (PDB) was employed. This includes: 1) a manually curated library of 77 X-ray crystal structures for RNA (Bernauer *et al.*, 2011), selected to be nonredundant and with resolution higher than 3.5 Å; 2) 175 crystal structures, selected from the dataset used in Svozil *et al.* (2008), containing non-complexed DNA structures adopting a wide variety of conformations (45 in A-form, 72 B-form, 39 Z-form and 19 quadruplexes). Hydrogens atoms were added with the Reduce software (Word *et al.*, 1999). All the listed structures were used for the calculation of the averaged form factors. For the validation we considered only a subset of these crystal structures, for which no missing non-hydrogen atoms neither missing or modified residues was present. This resulted in a validation set comprising 44 PDB structures for RNA and 121 for DNA. A complete list of the PDBs included in the datasets is given in Table S2.

### 2.5. Hybrid all-atom/coarse-grain SAXS calculation and PLUMED-ISDB implementation

The Martini form factors for both proteins (Niebling *et al.*, 2014) and nucleic acids (as computed in this work, see Table S3) are implemented in the PLUMED-ISDB (Bonomi & Camilloni, 2017) module and can be activated using the keyword “MARTINI” within the SAXS collective variable. An implementation of the atomic scattering factors, corrected by the excluded volume, can be activated with the “ATOMISTIC” keyword. It is also possible to define new form factors using a polynomial expansion of any order. This allows a high customizability of SAXS-driven simulations, which can be run in different modes: i. the atomistic mode, using both an all-atom force field and atomistic forward model; ii. the coarse-grain mode, using both Martini force field and form factor; iii. the hybrid multiresolution mode (described below) where the simulations are run with an atomistic force field and the Martini or other user-defined form factors are used for the forward model.

Importantly, to account for the approximations involved in the SAXS calculation (i.e. the coarse-grain representation does not consider the excess of electron density in the hydration shell, and the coarse graining itself), as well as for the noise in the data, we employed metainference (Bonomi *et al.*, 2016). Briefly, given a set of scattering vectors *q* and the measured intensities *I_q_*, if we assume that the global error can be modelled by a Gaussian per data point and that the measured and calculated intensities are defined modulo a multiplicative constant λ, one can show that an optimal balance between the force-field energy and the experimental data can be obtained by defining the metainference energy, *E_MI_*, as (Löhr *et al.*, 2017):

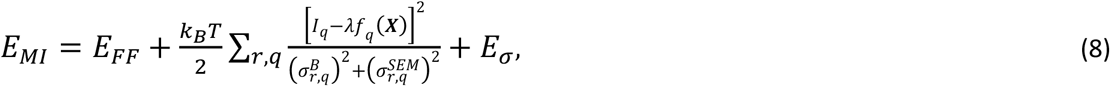

where *E_FF_* is the energy of the force field, *k_B_* the Boltzmann constant, *T* the temperature, *f_q_*(***X***) the calculated intensity (forward model) for the configuration ***X***, 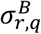 is an uncertainty parameter that describes random and systematic errors, 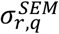 is the standard error of the mean related to the conformational averaging, and *E_σ_* is an energy term that accounts for normalization of the data likelihood and error priors. λ and 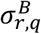 are sampled along with the MD by a Monte Carlo. The sum runs over the set of selected *q* and over multiple copies of the simulation. Importantly, if conformational averaging is not considered then the sum runs only over *q*, 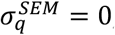, and metainference becomes equivalent to the Inferential Structure Determination approach (Rieping *et al*., 2005).

In the hybrid multi-resolution mode, given an atomic resolution structure, the position of Martini beads is calculated on-the-fly by PLUMED. The beads are associated to virtual atoms and subsequently are used in combination with the appropriate form factors to calculate the SAXS. As exemplified in the box, PLUMED first computes the coordinates of the centre of mass of each bead; the beads are then used by the SAXS action to calculate the Debye equation using the appropriate form factors.

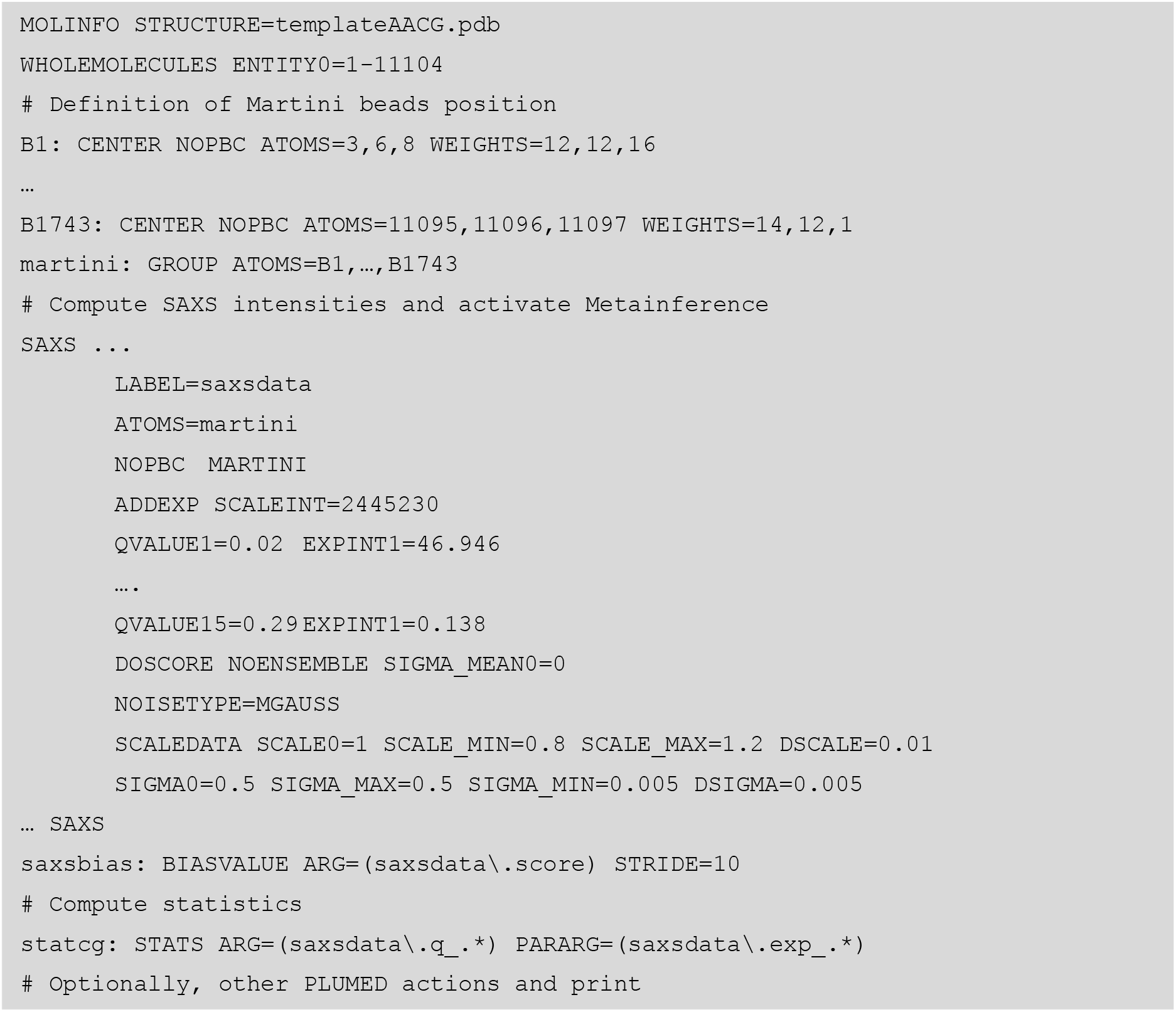

In order to correctly associate each bead to an atom type a PDB file must be provided (templateAACG.pdb in the example above), containing both the atomistic and the coarse-grain coordinates. Particular attention should be given to the atoms numbering, where the number of the first Martini bead should be equal to 1 + the number of atoms in the atomistic structure, comprising ions and solvent. The renumbering can be easily achieved using the PLUMED tool pdbrenumber, where numbers greater than 100000 are written in the hybrid 36 format.

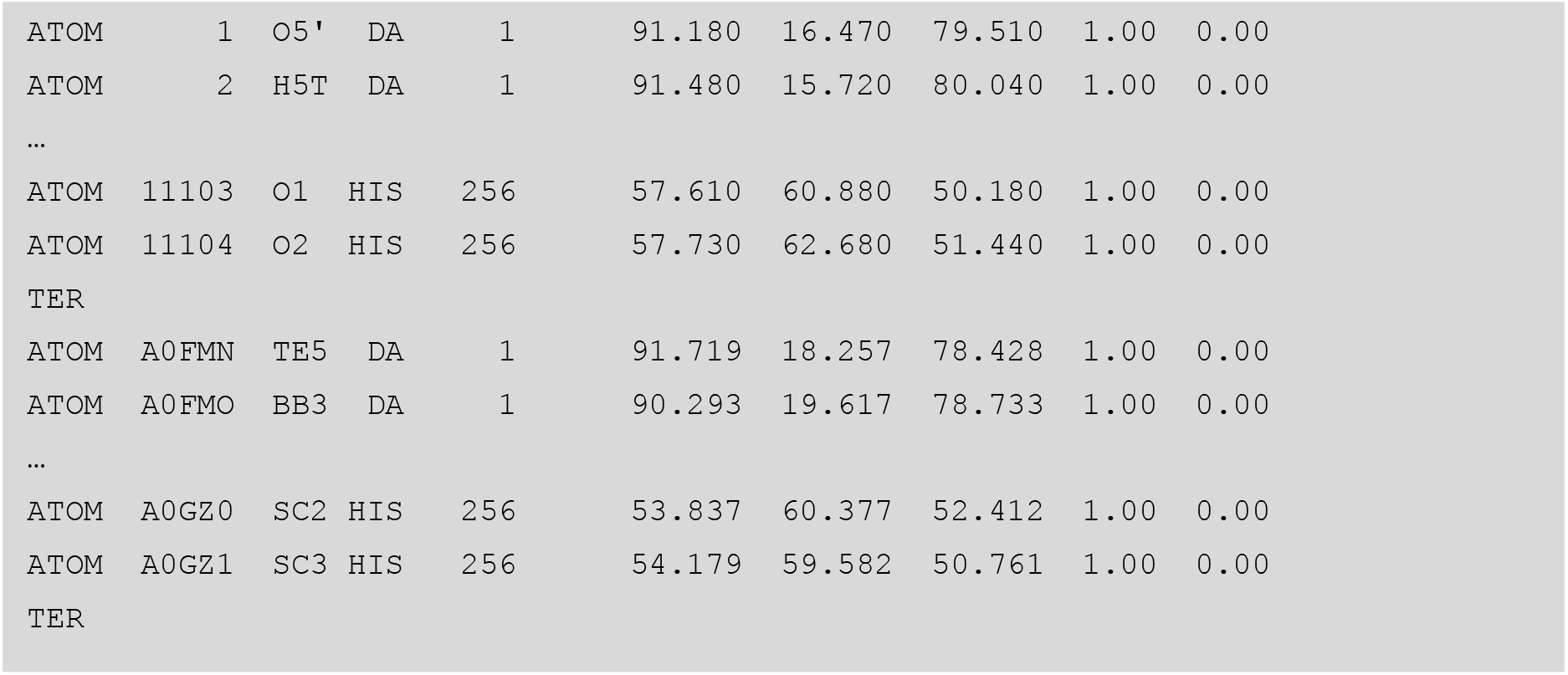

The SAXS results can be printed into an output file and, in case of a running MD simulation, can also be used in combination with METAINFERENCE (or other methods) to restrain the simulation. In the example metainference is activated by DOSCORE and the following keywords set the relevant parameters. The metainference energy is then applied using BIASVALUE every STRIDE step. It is worth noting that the flexibility of PLUMED allows to adopt a multiple time-step protocol for the integration of SAXS data in simulations, i.e. applying the metainference bias only at every few time steps (Ferrarotti *et al.*, 2015). This can be useful to further speed up the simulations and is fully justified in the case of SAXS data since the temporal fluctuations of this variable are slower than the ones in atomistic coordinates (Kimanius *et al.*, 2015).

Complete example files to run metainference simulations with the hybrid multi-resolution mode are provided in our GitHub repository https://github.com/carlocamilloni/papers-data.

### 2.6. Computational details of the simulations

MD simulations were performed on two protein/nucleic-acid complexes. First, the ComE-comcde DNA-protein complex, for which both experimental SAXS data and a calculated model are deposited in the SASBDB entry SASDAB7 (Sanchez et al., 2015; Valentini et al., 2015). Second, the complex of a single-stranded 12-mer oligonucleotide with a region of the heterogenous nuclear ribonucleoprotein A1 (Kooshapur et al., 2018), that we previously refined using standard atomistic scattering factors.

MD simulations were performed with GROMACS 2018, PLUMED 2 and the PLUMED-ISDB module (Tribello *et al.*, 2014; Abraham *et al.*, 2015; Bonomi & Camilloni, 2017). Both systems were prepared using the amber14sb force field for protein (Maier *et al.*, 2015) with parmbsc1 (Ivani *et al.*, 2015) and the TIP3P water model (Jorgensen *et al.*, 1983), solvated in a triclinic box and neutralized. After an initial energy minimization, the solute was equilibrated using the Berendsen thermostat (Berendsen *et al.*, 1984) to obtain the desired temperature of 300 K. For each system, one 5 ns production run was performed, in which metainference on a single replica was used to introduce SAXS restraints. For the protein/DNA complex an additional run without the inclusion of experimental information was performed as a reference. During the production runs, the md integrator was employed with a time step of 2 fs, temperature was controlled using the Bussi thermostat (Bussi *et al.*, 2007) and bonds were constrained with the LINCS algorithm (Hess *et al.*, 1998), using a matrix expansion on the order of 6 and 2 iterations per step. The van der Waals and short-range electrostatic interactions were truncated at 0.9 nm, whereas long-range electrostatic interactions were treated with the particle mesh Ewald method (Darden *et al.*, 1993).

In the case of the ComE-comcde DNA-protein complex, both the metainference and the unrestrained simulations were evolved for a total of 5 ns through a series of 20 simulated annealing cycles, with a period of 250 ps each and the temperature varying between 300 and 400 K. Specifically, each cycle consisted of 100 ps at 300 K, a fast increase of the temperature from 300 to 400 K, 20 ps at 400 K, and finally a linear cooling from 400 to 300 K in 120 ps. Only structures extracted from the intervals at 300 K in the last 10 cycles were used for analysis. In order to avoid the opening of DNA in the high temperature intervals, in both the simulations we restrained the hydrogen bonds between the first and last two couples of nucleotides adding a harmonic potential centred at 0.3 nm and with a force constant of 1000 kJ/(mol nm^2^). Specifically, the restraints were imposed on the distances between oxygens and nitrogens involved in hydrogen bonds for the couples A1-T76, A2-T75, A39-T39 and A37-T40. In the metainference simulation, a set of 15 representative SAXS intensities at different scattering vectors, ranging between 0.02 Å^−1^ and 0.3 Å^−1^, were also added as restraints. These representative intensities were extracted from the experimental data, where a 15-point running average was performed to reduce the influence of experimental noise. Metainference was applied every 10 steps, using a single Gaussian noise per data-point and sampling a scaling factor between experimental and calculated SAXS intensities with a flat prior between 0.8 and 1.2. An initial value for this scaling factor was chosen to match the experimental and calculated intensity at the scattering vector *q*=0.02 Å^−1^ for the initial model.

In the case of the RNA-protein complex, the metainference simulation was evolved for 5 ns maintaining the temperature at the value of 300 K. Restraints in the form of harmonic upper-wall potentials were applied as described in Kooshapur *et al.* (2018) to maintain critical protein-RNA interface contacts, salt bridges and protein secondary structures, as found in the related crystal structure (PDB: 6DCL). 43 representative SAXS intensities were used as restraints in metainference, corresponding to scattering vectors between 0.03 Å^−1^ and 0.45 Å^−1^. These intensities were obtained fitting experimental data with a 16^th^ degree polynomial up to scattering value of 0.5 Å^1^, following the work done in Kooshapur *et al.* (2018). Metainference was applied every 10 steps, using a single Gaussian noise per data-point and the scaling factor was sampled from a Gaussian prior.

For each run one reference model was selected clustering (based on the root mean square deviation of the position (RMSD)) the structures sampled at 300 K in the second half of the run and choosing the centre of the most populated cluster.

## 3. Results and discussion

### 3.1. Form Factors of DNA and RNA nucleotides

According to the Martini mapping scheme, the RNA and DNA nucleotides are represented with six or seven beads. Here we also considered two additional beads per nucleotide, representing the 5’ and 3’ terminal beads and being a simple modification (i.e. an addition of the terminal hydroxyl group) of the backbone beads BB2 and BB3. Indeed, while non-differentiating the terminal beads in protein is acceptable, this approximation in nucleic acids could result in computed scattering intensities with large deviations from the atomistic ones, especially when short oligomers are considered and for scattering vectors close to *q*=0. Totally, this results in the computation of 34 coarse-grain form factors for DNA and 34 for RNA, listed in Table S1.

For each bead, the effective form factor is calculated using equation (6) as described in the Methods section, averaging over a large number of different structures taken from the PDB (Table S2). The resulting coarse-grain form factors are represented in Figure 1 and listed in Table S3. We observed that beads with the same chemical composition in different nucleotides (i.e. all the backbone and terminal beads, purine beads SC1-SC4 and pyrimidine bead SC1) have perfectly superimposable Martini form factors. This holds true also when comparing corresponding beads in DNA and RNA, with the obvious exception of the SC3 bead for thymine/uracil and of the backbone BB3/TE3 beads, which in RNA contain the additional oxygen atom in position 2’. Of notice, the variability observed between the individual form factors to be averaged (Figure S1) is smaller than that observed in proteins (Niebling *et al.*, 2014). This can be explained by the highly conserved structural arrangement of the different atoms within each bead and it is a promising indication of reliability for coarse-grain scattering calculations. The only exceptions to this behaviour are represented by the terminal beads, which display larger variation due to the different orientations that the terminal hydroxyl group can assume with respect to the other atoms of the bead.

**Figure 1.**
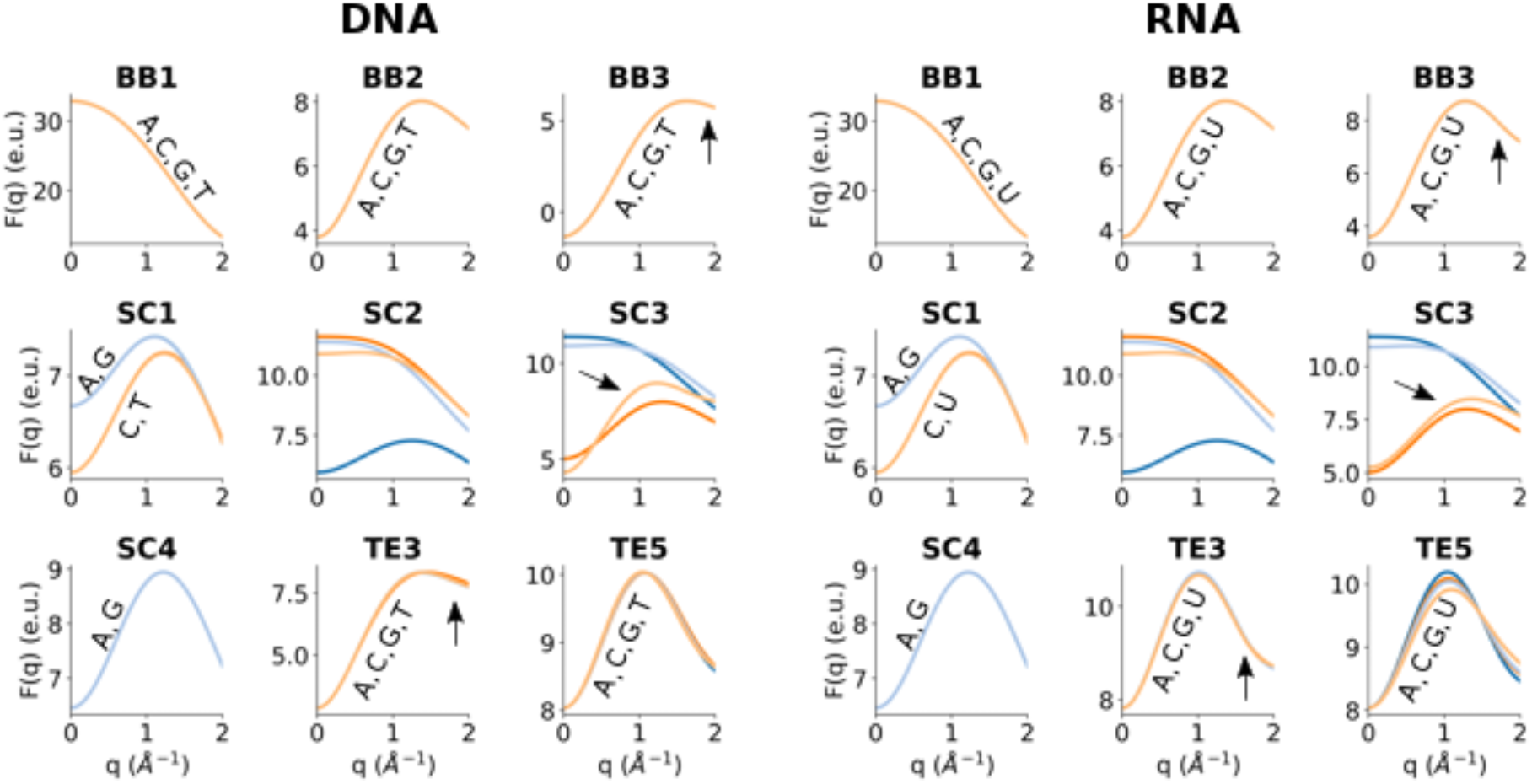
Calculated coarse-grain form factors for DNA (left) and RNA (right). The form factors are represented for all Martini beads of the nucleotides: adenine (blue), cytosine (dark orange), guanine (light blue) and thymine/uracil (light orange). When two or more structure factures are superimposed, labels are added for clarity. Differences between corresponding DNA and RNA beads are highlighted with black arrows.

### 3.2. Comparison of scattering intensities computed with all-atom or coarse-grain form factors

To evaluate the accuracy of Martini form factors in computing scattering intensities for nucleic acids, we compared coarse-grain SAXS profiles with the atomistic ones for a library of crystal structures, comprising 44 RNA and 121 DNA structures (Table S2). The differences between Martini and atomistic curves have been measured calculating the average relative squared error over different *q*-values:

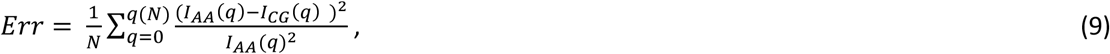

where *I_AA/CG_* are the intensities computed with all-atom or Martini form factors, respectively, *N* is the number of data points and *q* varies between 0 and *q*(*N*), being the step between *q*(*i*) and *q*(*i-1*) equal to 0.01 Å^−1^. To compare our results with the ones obtained for protein in Niebling *et al.* (2014), we identified the maximum *q* value for which the error of equation (9) is smaller than 0.2%. This value, denoted as *q_threshold_*, has been computed for each of the considered RNA and DNA structures and its distribution is reported in Figure 2. The average *q_threshold_* values for RNA and DNA (0.47 and 0.88 Å^−1^, respectively) are comparable with the one previously found for protein (0.53 Å^−1^) and suggest that the scattering intensities up to *q*~0.45 Å^−1^ can be reliably calculated using the coarse-grain approximation. Figure 2 highlights a distinct behaviour for DNA and RNA: while the *q_threshold_* values display small deviations in the RNA structures, they are considerably spread in the case of DNA. In particular, we found that different DNA conformations are associated with diverse values of *q_threshold_*, where DNA in A- or Z-form mainly display *q_threshold_* between 0.4 and 0.6 Å^−1^, while B-form DNA structures often reach values greater than 1.0 Å^−1^, conceivably due to their less compact structure (see Figure S2).

**Figure 2.**
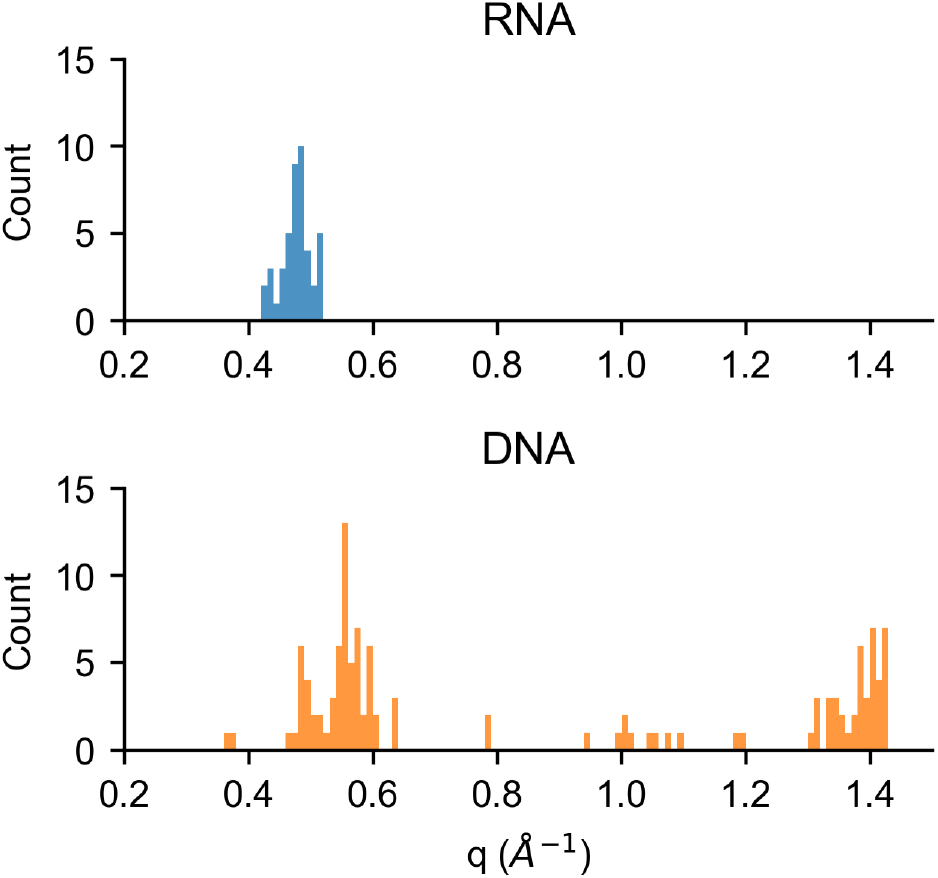
Distribution of *q_threshold_* values for 44 RNA and 121 DNA crystallographic structures. Each *q_threshold_* has been computed comparing atomistic and coarse-grain scattering curves and represents the maximum *q* values for which the error defined in equation (8) is smaller than 0.2%.

As an additional metric to assess the accuracy of coarse-grain intensities computation, we adopted the *R*-scoring function:

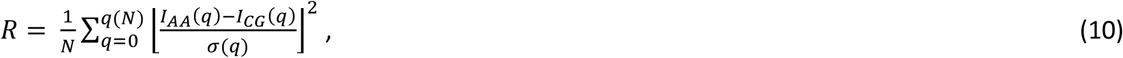

where *σ*(*q*) = *I*(*q*)(*q* + *a*) *b*, with *a* = 0.15 and *b* = 0.3 (Tong *et al.*, 2016; Stovgaard *et al.*, 2010). This value aims to reproduce the usual *χ*^2^ metric in evaluating differences between SAXS profiles, where an empirical standard deviation is adopted since experimental errors are not available in theoretical curves. The form of *σ*(*q*) and the values of the *a* and *b* parameters were chosen as in Stovgaard *et al.* (2010), to be stricter in the portion of the curve of major interest for structure prediction (*q* values lower than 0.5 Å^−1^). In Figure 3, the distribution of *R*-values for RNA and DNA evaluated for *q* up to 0.5 Å^−1^ is reported, along with the average *R*-value as a function of the scattering vector *q* used as cut-off. For DNA, 99% of the structures present an *R*-value lower than 0.1, with intensities for DNA in B-form being reproduced again slightly better than for the other forms (Figure S3). For RNA, only 68% of the structures display an *R*-value below 0.1 as a consequence of a sharp increase of *R* for scattering vectors between 0.4 and 0.5 Å^−1^. By decreasing the *q* cut-off to 0.45 Å^−1^ we found that 95% of the structures satisfies *R*<0.1 (Figure S4), further confirming this range as optimal for coarse-grain intensities calculations involving RNA molecules.

**Figure 3.**
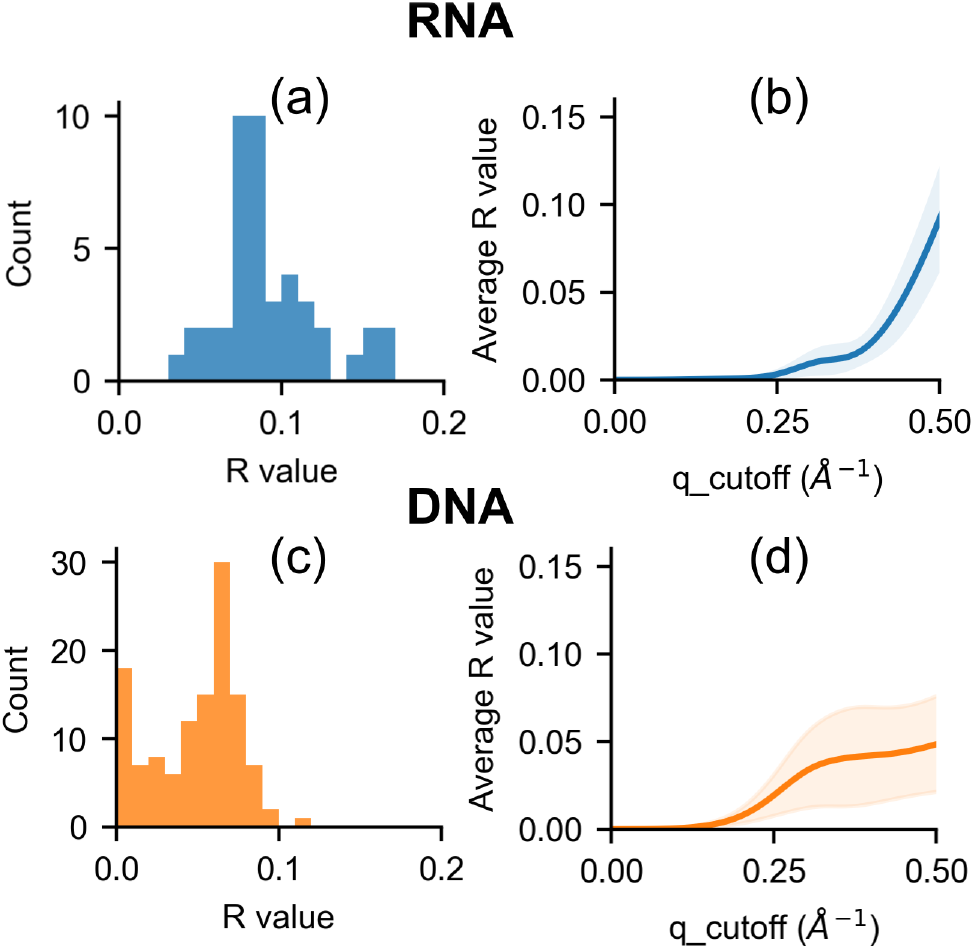
Distribution of *R*-values for 44 RNA (a) and 121 DNA (c) crystallographic structures, computed over a range of scattering vector *q* below 0.5 Å^−1^. *R*-values, averaged over the whole set of structures for RNA (b) and DNA (d) and evaluated over a range of scattering vectors below a cut-off (*q_cutoff*), are reported as a function of the cut-off. The standard deviation is represented as a shadow.

The form factors herein derived for nucleic acids can be seamlessly combined with those for proteins and then used for an efficient back-calculation of SAXS curves in protein/nucleic-acid complexes for scattering vectors up to 0.45 Å^−1^. The computational efficiency gained by the use of these coarse-grain form factors is very important in applications where repeated evaluations of scattering intensities are requested. In particular, they could be exploited to drive MD simulations to match SAXS data, allowing an extension of the system size and the simulation length with respect to previous applications. The Martini form factors can be naturally included in simulations run with the coarse-grain Martini force field. Moreover, we propose a hybrid coarse-grain/all-atom approach, where the simulations are run with full atomistic details, while the Martini form factors are used for the SAXS calculation, thus allowing a faster back-calculation of the scattering intensities (cf. Theory and Methods).

### 3.3. Refinement of protein/nucleic acid complexes against SAXS data

To demonstrate the efficiency and reasonable accuracy of the Martini form factors in experimental driven MD simulations, we exploited them in the refinement of protein/nucleic-acid complexes against SAXS data. To this aim we took advantages of the metainference technique, which allows the introduction of noisy and ensemble-averaged experimental data in MD simulations. Importantly, metainference also takes errors resulting from the forward model into account. This is particularly relevant here since the approximations resulting from the coarse-grain representation do not consider the excess of electron density in the hydration shell.

#### 3.3.1. ComE-comcde DNA-protein complex

ComE is a two-domain protein, part of the ComD-ComE two-component signalling system, which dimerizes in solution via its REC domain when activated by ComD-induced phosphorylation. In Sanchez et al. (2015), SAXS data were used to show that the ComE^D58E^ active mimic mutant is found in dimeric form when bound to the promoter region comcde. Furthermore, it induces an extra-bending of DNA. In that work, a model, comprising of two ComE bound to the 38-mer comcde duplex, was build to fit with SAXS data exploiting the available crystallographic structure of the ComE dimer (PDB: 4CBV, (Boudes *et al.*, 2014)). The model proposed by Sanchez and coworkers displays a good agreement with SAXS data and provides interesting structural insights into ComE-comcde binding mode. Here, we show that our metainference-based approach could be exploited to improve the quality of the model, solving steric clashes and other defects created during the modelling phase, meanwhile further improving the agreement with SAXS data. After a short energy minimization of the system, whose initial coordinates were taken from the SASBDB entry SASDAB7, we performed a 5 ns long simulated annealing (SA) simulation in which metainference was used to introduce SAXS restraints (see Theory and Methods section). We selected the refined model clustering the structures sampled at 300K in the last 10 SA cycles (out of 20) based on geometrical similarity and choosing the centre of the most populated cluster. To check the importance of using SAXS data, we also performed an additional run using the same SA protocol without metainference (i.e. without introducing experimental restraints). The agreement with SAXS data and model quality were assessed using *CRYSOL* and Molprobity (Adams *et al.*, 2010; Davis *et al.*, 2007) comparing 4 different structures: the initial model, the initial model after energy minimization, the refined structure extracted from the metainference simulation and the structure selected from the unrestrained simulation. While the energy minimization can solve most of the steric clashes, metainference simulations are useful to further improve both the agreement with experimental data and the quality of the model in terms of Ramachandran and clash-score (Table 1). The refined model (Figure 4) displays a *χ*^2^ of 1.19, slightly better compared to the initial one, maintains the known critical interactions between ComE and the DNA recognition sites (mainly involving residues H168, K203 and K235) and shows only small deviations from the crystal structures of the ComE dimer (backbone-RMSD of 2.5 Å with respect to the reference structure in PDB 4CBV). Conversely, the structure extracted from the unrestrained simulation, even if having a good Molprobity score, misses most of the DNA-ComE contacts and significantly alter the ComE dimer conformation, showing a backbone-RMSD of 6.3 Å with respect to the crystal structure. Overall, this results in a poor agreement with experimental data, confirmed by a *χ*^2^ > 3. Of notice, all the *χ*^2^ values reporting the distance from experimental SAXS data were calculated with *CRYSOL* and are therefore independent from the strategy used to back-calculate SAXS intensities and consequently restrain the simulations.

**Table 1.**
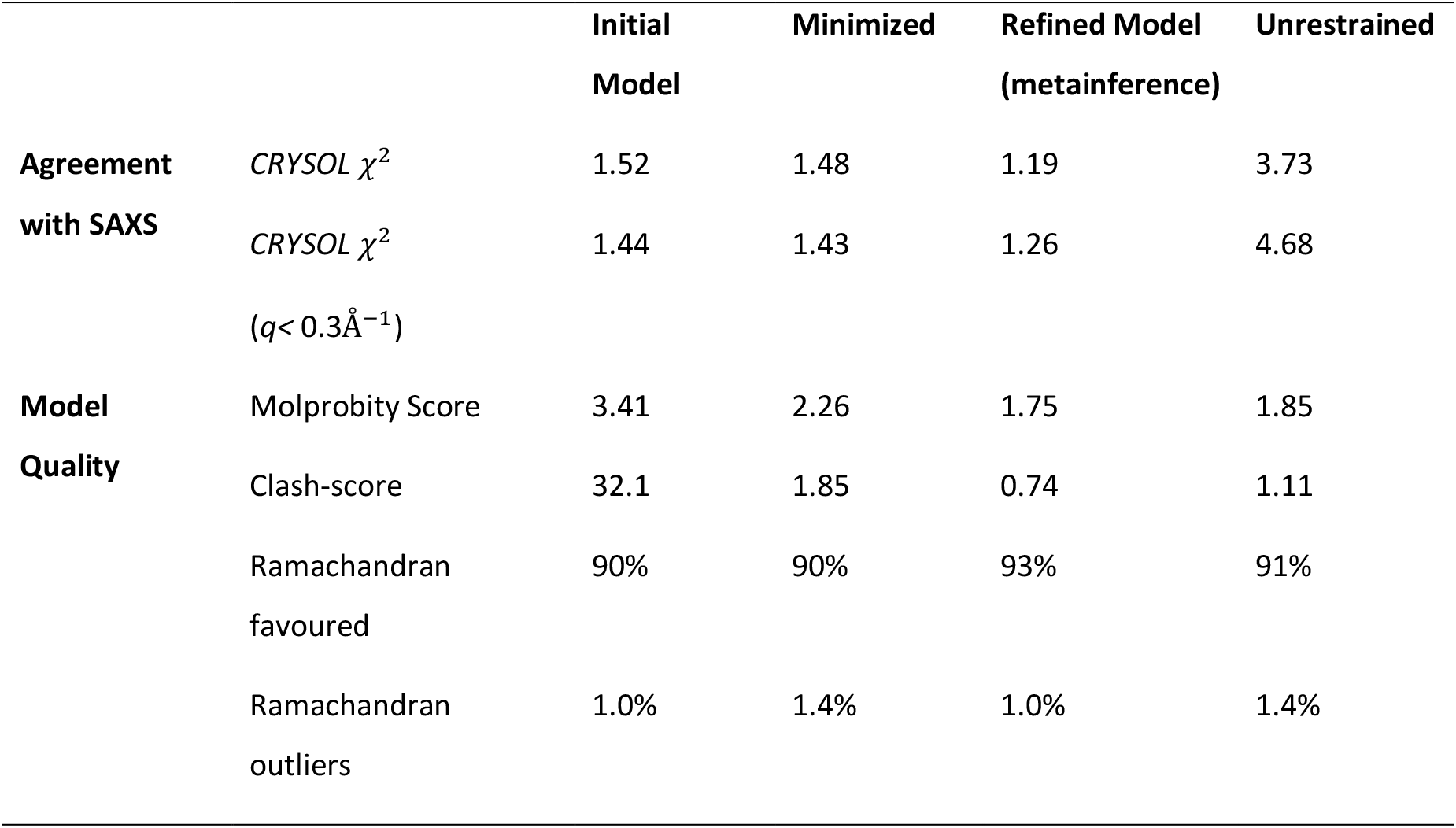
Evaluation of representative protein/DNA structures in terms of agreement with SAXS data and model quality. The structures considered are: i. the initial model; ii. the initial model after energy minimization; iii. a refined model extracted from metainference simulation; iv. a representative structure extracted from the unrestrained simulation. The agreement with SAXS data was measured with *CRYSOL* (Svergun *et al.*, 1995) using the maximum order of harmonics available and 18 points for the Fibonacci grid. The model quality was assessed using the Molprobity validation implemented in Phenix (Adams *et al.*, 2010; Davis *et al.*, 2007).

**Figure 4.**
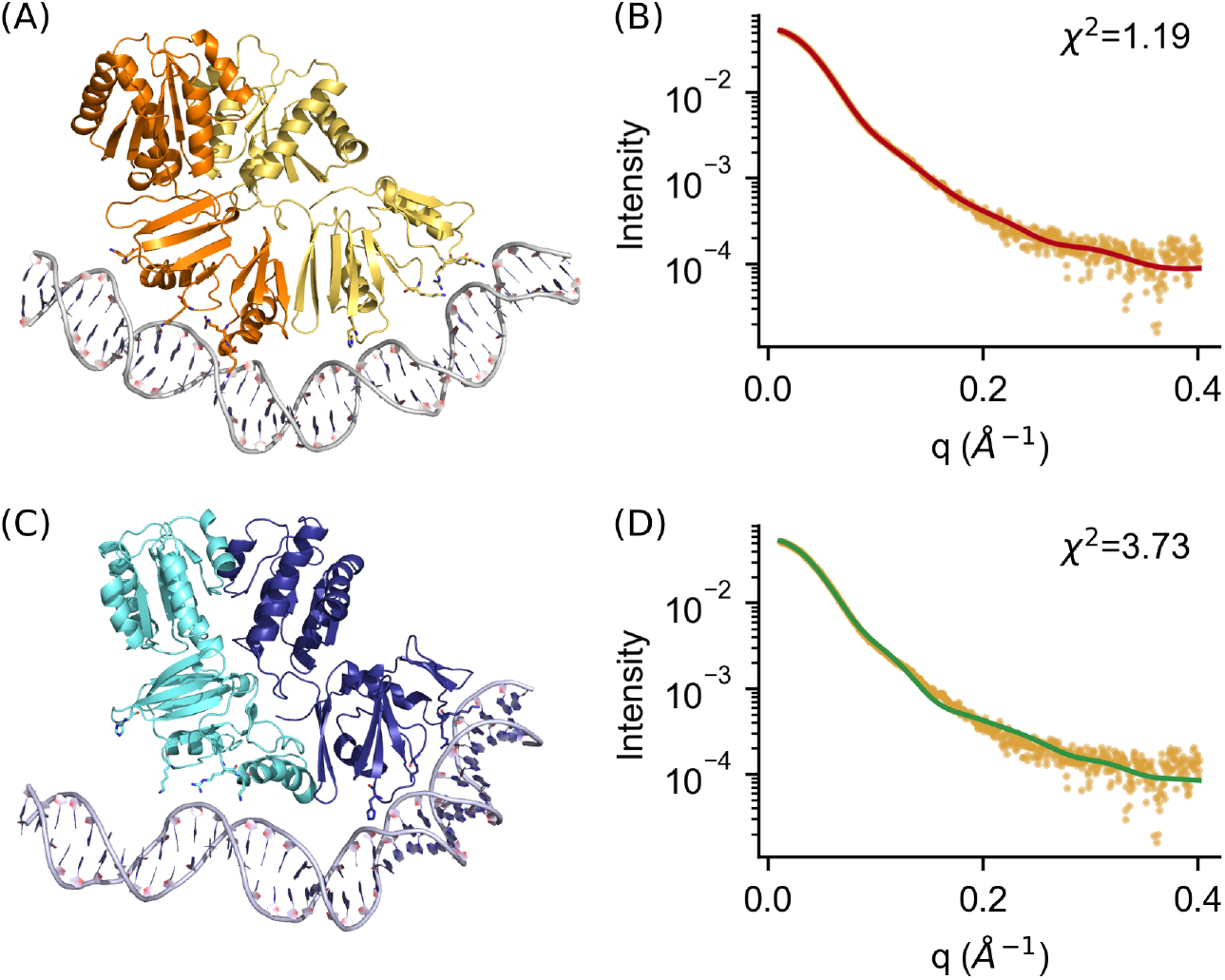
The representative ComE-DNA structures extracted from the metainference (A, B) and unrestrained (C, D) simulations are shown, along with their fitting to experimental data, represented with yellow points, according to *CRYSOL*.

Importantly, we observed that the integration of SAXS data in simulation is prohibitive if atomistic scattering factors are used for the back-calculations of the intensities; however, its impact on MD performances is significantly reduced exploiting the Martini form factors and adopting the multiple time-steps strategy, where the metainference bias is applied only every 10 time-steps (Table 2).

**Table 2.**
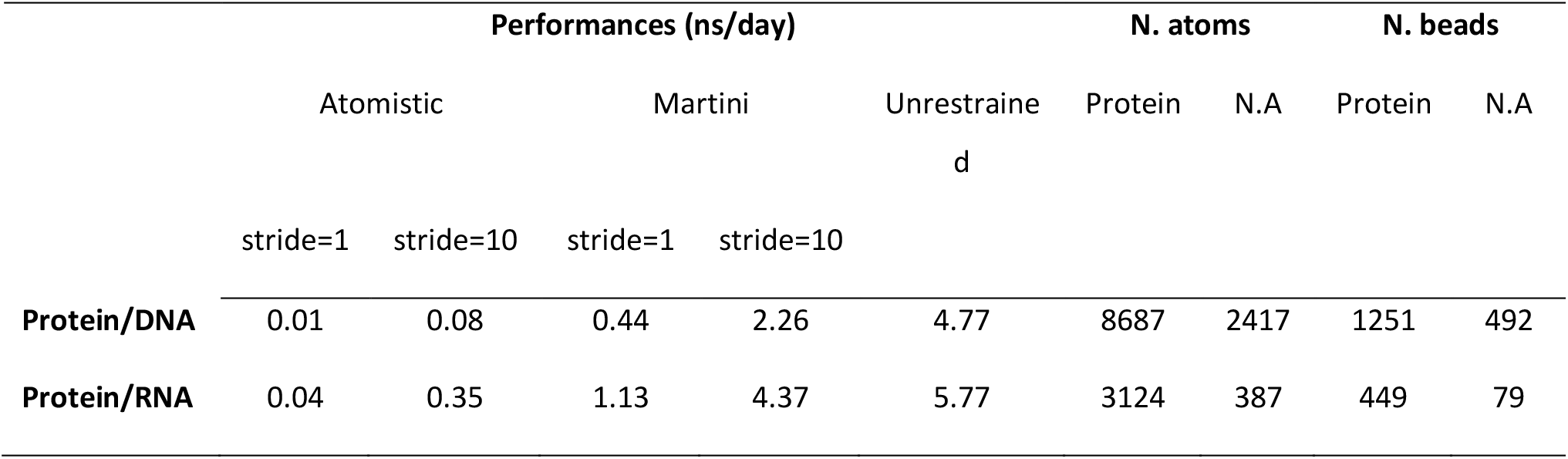
Summary of the performances for protein/DNA and protein/RNA simulations, achieved using the metainference approach (computing the scattering intensities with either atomistic or Martini form factors) or without the integration of SAXS data (unrestrained). Data for metainference simulations in which the bias is applied every 10 time-steps are reported. Performances were estimated on an Intel Xeon E5 3.5 GHz using 4 cores. The number of protein/nucleic acid atoms and beads for each system is also indicated.

#### 3.3.2. The UP1-miRNA complex

As a second test system we used a protein-RNA complex that we previously refined against SAXS and NMR data (Kooshapur *et al.*, 2018). This complex involves the binding of a single-stranded 12-mer oligonucleotide, derived from the micro RNA 18a primary transcript, and the Unwinding Protein 1 (UP1), comprising two tandem domains that constitute the RNA-binding region of the heterogenous nuclear ribonucleoprotein A1. The crystal structure of UP1/12-mer RNA previously solved (PDB 6DCL) showed a 2:2 stoichiometry that was found to be not representative of the 1:1 stoichiometry measured in solution. Therefore, a refinement against experimental data in solution was performed, where the key features of the crystallographic binding interface were retained leading to a model of UP1/12-mer RNA with the correct 1:1 stoichiometry (Kooshapur *et al.*, 2018). Here we reproduce this same refinement procedure, using an analogous approach where the main protein/RNA and protein/protein interaction sites are restrained (see Theory and Methods). The resulting UP1/12-mer RNA model, extracted from our metainference simulation as described in the Method section, is analogous to the reference one (obtained via a fully-atomistic approach) both in terms of agreement with SAXS data and model quality (Table S4). The representative metainference-derived model and the fitting with experimental data, according to *CRYSOL*, are shown in Figure 5. Of note, the adoption of Martini form factors in combination with a multiple time-step scheme allows to approach the performance of the unrestrained simulation (4.4 ns/day vs 5.8 ns/day), outperforming the simulations relying on atomistic scattering factors (0.35 ns/day also when applying metainference bias every 10 time-steps, see Table 2).

**Figure 5.**
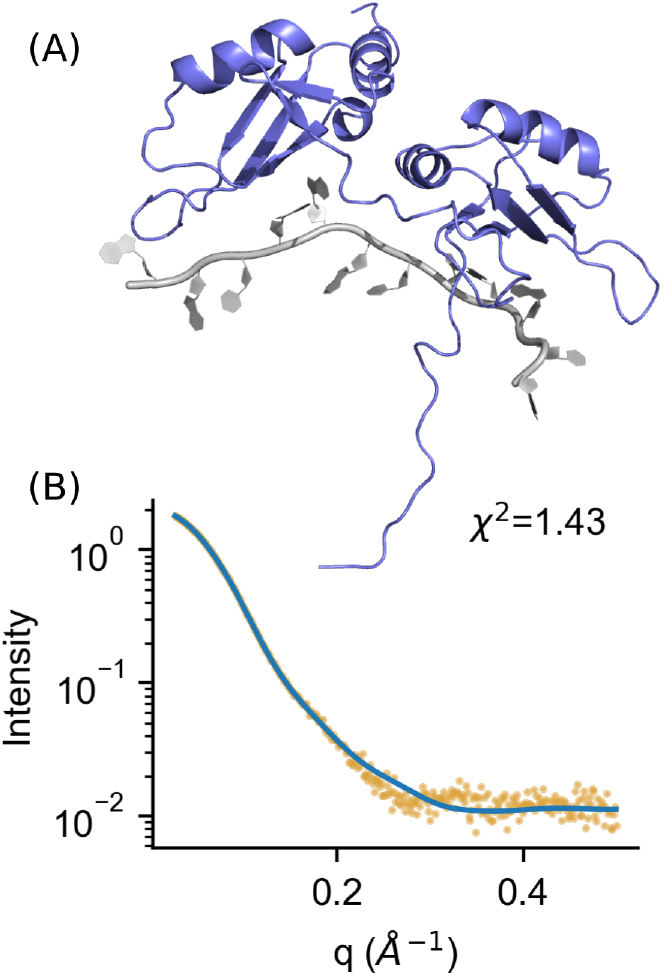
A representative UP1-miRNA structure extracted from metainference (A) is shown, along with its fitting to experimental data according to *CRYSOL* (B).

## 4. Conclusion

The cost of computing scattering intensities from atomic structures is a limiting factor for the integration of SAXS experimental data in MD simulations and for other applications where multiple evaluations of scattering curves are required. Here we extended the work of Niebling *et al.* (2014) to nucleic acids, computing Martini beads form factors for RNA and DNA and showing that they can be exploited to accurately evaluate SAXS intensities for scattering vectors up to 0.45 Å^−1^. Further, we implemented these coarse-grain form factors in PLUMED and showed how they could be used for the structure refinement of molecular systems against SAXS data adopting a hybrid atomistic/coarse-grain approach. Overall, our results clearly indicate that Martini form factors, for both proteins and nucleic acids, can be safely used to restrain atomistic simulations against SAXS intensities, reproducing experimental data with an accuracy comparable to the one achieved in atomistic mode and improving the performances up to a factor of 50. We anticipate that our protocol, by using metainference, can be used in combination with other experimental data, as well as extended to run multiple-replica simulations taking then into account molecular conformational averaging. Lastly, we note that the applicability of the Martini form factors is not limited to their use in SAXS-driven MD simulations, but they could also be used for the analysis of single structures or trajectories exploiting the PLUMED driver utility.

## Supporting information

## Acknowledgements

We acknowledge the CINECA award under the ISCRA initiative, for the availability of high-performance computing resources and support.

