## Supplementary material for "Martini bead form factors for nucleic-acids and their application in the refinement of protein/nucleic-acid complexes against SAXS data"

### Supporting information

**Table S1** List of the scattering types used and their atomic composition. The sum of the reduced atomic scattering factor at $q = 0$ (i.e. $F_i(q = 0) = \sum_{k \in i} f'_k$) is also reported. The cases in which this sum results in a negative value, and thus need a correction as described in the text, are highlighted in bold.

| Nucleotide | Bead | Number of atoms | | | | | $F(q = 0)$ |
|---|---|---|---|---|---|---|---|
| | | C | H | N | O | P | |
| DA | BB1 | 0 | 0 | 0 | 4 | 1 | 32.89 |
| DA | BB2 | 2 | 3 | 0 | 1 | 0 | 3.81 |
| DA | BB3 | 3 | 4 | 0 | 0 | 0 | **-1.36** |
| DA | SC1 | 1 | 0 | 1 | 0 | 0 | 6.67 |
| DA | SC2 | 1 | 1 | 1 | 0 | 0 | 5.95 |
| DA | SC3 | 1 | 2 | 2 | 0 | 0 | 11.39 |
| DA | SC4 | 2 | 1 | 1 | 0 | 0 | 6.46 |
| DA | TE3 | 3 | 5 | 0 | 1 | 0 | 2.87 |
| DA | TE5 | 2 | 4 | 0 | 2 | 0 | 8.04 |
| DC | BB1 | 0 | 0 | 0 | 4 | 1 | 32.89 |
| DC | BB2 | 2 | 3 | 0 | 1 | 0 | 3.81 |
| DC | BB3 | 3 | 4 | 0 | 0 | 0 | **-1.36** |
| DC | SC1 | 1 | 1 | 1 | 0 | 0 | 5.95 |
| DC | SC2 | 1 | 0 | 1 | 1 | 0 | 11.62 |
| DC | SC3 | 2 | 3 | 1 | 0 | 0 | 5.02 |
| DC | TE3 | 3 | 5 | 0 | 1 | 0 | 2.87 |
| DC | TE5 | 2 | 4 | 0 | 2 | 0 | 8.04 |
| DG | BB1 | 0 | 0 | 0 | 4 | 1 | 32.89 |
| DG | BB2 | 2 | 3 | 0 | 1 | 0 | 3.81 |
| DG | BB3 | 3 | 4 | 0 | 0 | 0 | **-1.36** |
| DG | SC1 | 1 | 0 | 1 | 0 | 0 | 6.67 |
| DG | SC2 | 1 | 2 | 2 | 0 | 0 | 11.39 |
| DG | SC3 | 1 | 1 | 1 | 1 | 0 | 10.90 |
| DG | SC4 | 2 | 1 | 1 | 0 | 0 | 6.46 |
| DG | TE3 | 3 | 5 | 0 | 1 | 0 | 2.87 |
| DG | TE5 | 2 | 4 | 0 | 2 | 0 | 8.04 |
| DT | BB1 | 0 | 0 | 0 | 4 | 1 | 32.89 |
| DT | BB2 | 2 | 3 | 0 | 1 | 0 | 3.81 |
| DT | BB3 | 3 | 4 | 0 | 0 | 0 | **-1.36** |
| DT | SC1 | 1 | 1 | 1 | 0 | 0 | 5.95 |
| DT | SC2 | 1 | 1 | 1 | 1 | 0 | 10.90 |
| DT | SC3 | 3 | 3 | 0 | 1 | 0 | 4.31 |
| DT | TE3 | 3 | 5 | 0 | 1 | 0 | 2.87 |
| DT | TE5 | 2 | 4 | 0 | 2 | 0 | 8.04 |
| A | BB1 | 0 | 0 | 0 | 4 | 1 | 32.89 |
| A | BB2 | 2 | 3 | 0 | 1 | 0 | 3.81 |
| A | BB3 | 3 | 4 | 0 | 1 | 0 | 3.59 |
| A | SC1 | 1 | 0 | 1 | 0 | 0 | 6.67 |
| A | SC2 | 1 | 1 | 1 | 0 | 0 | 5.95 |
| A | SC3 | 1 | 2 | 2 | 0 | 0 | 11.39 |
| A | SC4 | 2 | 1 | 1 | 0 | 0 | 6.46 |
| A | TE3 | 3 | 5 | 0 | 2 | 0 | 7.82 |
| A | TE5 | 2 | 4 | 0 | 2 | 0 | 8.04 |
| C | BB1 | 0 | 0 | 0 | 4 | 1 | 32.89 |
| C | BB2 | 2 | 3 | 0 | 1 | 0 | 3.81 |
| C | BB3 | 3 | 4 | 0 | 1 | 0 | 3.59 |

IMPORTANT: this document contains embedded data - to preserve data integrity, please ensure where possible that the IUCr Word tools (available from http://journals.iucr.org/services/docxtemplate/) are installed when editing this document.

| | | | | | | | |
|---|---|---|---|---|---|---|---|
| C | SC1 | 1 | 1 | 1 | 0 | 0 | 5.95 |
| C | SC2 | 1 | 0 | 1 | 1 | 0 | 11.62 |
| C | SC3 | 2 | 3 | 1 | 0 | 0 | 5.02 |
| C | TE3 | 3 | 5 | 0 | 2 | 0 | 7.82 |
| C | TE5 | 2 | 4 | 0 | 2 | 0 | 8.04 |
| G | BB1 | 0 | 0 | 0 | 4 | 1 | 32.89 |
| G | BB2 | 2 | 3 | 0 | 1 | 0 | 3.81 |
| G | BB3 | 3 | 4 | 0 | 1 | 0 | 3.59 |
| G | SC1 | 1 | 0 | 1 | 0 | 0 | 6.67 |
| G | SC2 | 1 | 2 | 2 | 0 | 0 | 11.39 |
| G | SC3 | 1 | 1 | 1 | 1 | 0 | 10.90 |
| G | SC4 | 2 | 1 | 1 | 0 | 0 | 6.46 |
| G | TE3 | 3 | 5 | 0 | 2 | 0 | 7.82 |
| G | TE5 | 2 | 4 | 0 | 2 | 0 | 8.04 |
| U | BB1 | 0 | 0 | 0 | 4 | 1 | 32.89 |
| U | BB2 | 2 | 3 | 0 | 1 | 0 | 3.81 |
| U | BB3 | 3 | 4 | 0 | 1 | 0 | 3.59 |
| U | SC1 | 1 | 1 | 1 | 0 | 0 | 5.95 |
| U | SC2 | 1 | 1 | 1 | 1 | 0 | 10.90 |
| U | SC3 | 2 | 1 | 0 | 1 | 0 | 5.25 |
| U | TE3 | 3 | 5 | 0 | 2 | 0 | 7.82 |
| U | TE5 | 2 | 4 | 0 | 2 | 0 | 8.04 |

**Table S2** PDB codes used for the calculations of the coarse-grained form factors, listed by category. The PDBs used in the validation are highlighted with a star. The numbers in parenthesis indicate the total number of structures in the category and the number of PDBs used for the validation, respectively.

| | | |
|---|---|---|
| RNA (77,44) | | 1z58, 357d, 1t0e*, 1t0d, 2oe6*, 1q93*, 1msy*, 1sdr*, 353d*, 361d*, 1dqf*, 1mme*, 2a64*, 2a2e, 1y0q*, 1x8w*, 1u9s*, 1nbs*, 2h0s, 1x9k*, 1x9c, 1ykq, 1nuj*, 1fir, 1i9v, 1yfg, 2tra, 434d*, 2il9*, 1kh6, 1xjr, 1k9w*, 2b8r*, 2nok*, 1csl*, 1l2x, 397d*, 1duq*, 1jzv, 1duh, 1d4r, 1z43*, 1kxk*, 1i9x*, 1mhk*, 387d*, 1yzd, 1f1t, 1kfo, 406d, 405d*, 1f27, 1qbp, 433d*, 413d*, 157d*, 205d*, 255d*, 280d*, 1kd5*, 1p79, 2ao5, 1j9h, 438d*, 2g91, 1g2j, 1sa9*, 259d*, 2a0p, 2g3s*, 333d, 402d*, 409d, 472d*, 1l3z*, 377d |
| DNA (175,121) | A-form (45,36) | 118d*, 137d*, 138d*, 160d*, 1d78*, 1d79*, 1dnz*, 1kgk, 1m77*, 1ma8, 1mlx, 1nzg, 1vj4*, 1xjx*, 1z7i, 1zex*, 1zey*, 1zf1*, 1zf6*, 1zf8*, 1zf9*, 1zfa*, 213d, 243d*, 260d*, 295d*, 2d94*, 317d*, 338d, 344d, 345d, 348d*, 349d*, 368d*, 369d*, 370d*, 371d*, 395d*, 396d*, 399d*, 414d*, 440d*, 9dna*, 1vt5*, 1vtb* |
| | B-form (72,51) | 122d, 123d, 158d*, 183d, 196d*, 1bd1*, 1bna*, 1cw9, 1d23*, 1d3r, 1d49*, 1d56*, 1d61, 1d8g*, 1d8x*, 1dou*, 1dpn, 1edr, 1ehv*, 1en3*, 1en8*, 1en9*, 1ene*, 1enn*, 1fq2*, 1g75, 1i3t, 1ikk*, 1j8l, 1jgr*, 1l4j*, 1l6b, 1m6g*, 1n1o, 1nvn*, 1nvy*, 1p4y*, 1p54, 1s23*, 1s2r*, 1sgs*, 1sk5*, 1ub8*, 1ve8, 1zf0*, 1zf3*, 1zf4*, 1zf5*, 1zf7*, 1zfb*, 1zff*, 1zfg*, 232d*, 251d*, 2d25, 307d*, 355d*, 3dnb*, 403d, 423d*, 428d*, 431d*, 436d, 454d, 455d*, 456d, 460d, 463d*, 476d*, 477d*, 5dnb*, 9bna* |
| | Z-form (39,21) | 131d*, 145d, 181d*, 1d40, 1d41, 1d48*, 1d53*, 1d76, 1da2, 1dcg*, 1dj6*, 1dn4, 1dn5, 1dnf, 1i0t*, 1ick*, 1jes, 1ljx*, 1omk, 1xa2*, 1xam*, 1zna*, 210d, 211d, 242d, 292d*, 293d*, 2dcg*, 313d, 314d*, 331d*, 336d*, 351d*, 362d*, 400d |
| | Quadruplexes (19,13) | 184d*, 190d*, 191d*, 1bqj*, 1cn0*, 1jpq, 1l1h*, 1mf5*, 1o0k, 1qyk*, 1qyl*, 1v3n, 1v3o, 1v3p, 200d*, 241d*, 244d*, 284d*, 352d |

**Table S3** List of the coefficients, to be used in a polynomial expansion of the sixth order, for each nucleotide bead.

| Nucl | Bead | A0 | A1 | A2 | A3 | A4 | A5 | A6 |
|---|---|---|---|---|---|---|---|---|
| DA | BB1 | 32.885000 | 0.081799 | -7.317359 | 2.156145 | -3.522632 | 2.306047 | -0.392701 |
| DA | BB2 | 3.806000 | -0.105977 | 9.525375 | -6.129910 | -0.540926 | 1.154291 | -0.215035 |
| DA | BB3 | -1.356000 | 0.589283 | 6.718941 | 4.140509 | -9.658599 | 4.431850 | -0.646573 |
| DA | SC1 | 6.671000 | -0.008714 | 1.632891 | -0.066377 | -1.486329 | 0.785518 | -0.120873 |
| DA | SC2 | 5.951000 | -0.026343 | 2.548643 | -0.490158 | -1.553869 | 0.866302 | -0.135462 |
| DA | SC3 | 11.394000 | 0.008595 | -0.254714 | 0.487188 | -1.745200 | 0.992462 | -0.163519 |
| DA | SC4 | 6.459000 | 0.019918 | 4.179623 | 0.974691 | -5.029504 | 2.553718 | -0.391134 |
| DA | TE3 | 2.874000 | 0.001129 | 12.511672 | -7.675480 | -2.022340 | 2.508371 | -0.494585 |
| DA | TE5 | 8.036000 | 0.004731 | 4.655544 | 0.664241 | -6.621313 | 3.961074 | -0.690758 |
| DC | BB1 | 32.885000 | 0.081899 | -7.324935 | 2.159769 | -3.526121 | 2.310586 | -0.394027 |
| DC | BB2 | 3.806000 | -0.105598 | 9.525277 | -6.121317 | -0.548994 | 1.155929 | -0.214945 |
| DC | BB3 | -1.356000 | 0.555257 | 6.803055 | 4.059247 | -9.610347 | 4.412538 | -0.643151 |
| DC | SC1 | 5.951000 | -0.028999 | 2.595878 | -0.553883 | -1.563951 | 0.889674 | -0.140625 |
| DC | SC2 | 11.621000 | 0.013581 | -0.249130 | 0.487872 | -1.528673 | 0.836949 | -0.133953 |
| DC | SC3 | 5.019000 | -0.032984 | 5.542428 | -0.960815 | -3.710516 | 2.165002 | -0.350234 |
| DC | TE3 | 2.874000 | -0.052355 | 13.092012 | -9.481282 | -0.149586 | 1.755372 | -0.393475 |
| DC | TE5 | 8.036000 | -0.005136 | 4.677057 | 0.483333 | -6.345110 | 3.833885 | -0.673678 |
| DG | BB1 | 32.885000 | 0.081829 | -7.321339 | 2.157679 | -3.523697 | 2.308396 | -0.393483 |
| DG | BB2 | 3.806000 | -0.106181 | 9.541690 | -6.151776 | -0.534624 | 1.155813 | -0.215670 |
| DG | BB3 | -1.356000 | 0.574891 | 6.751647 | 4.113009 | -9.633946 | 4.416754 | -0.643399 |
| DG | SC1 | 6.671000 | -0.008866 | 1.633330 | -0.068921 | -1.486835 | 0.786708 | -0.121139 |
| DG | SC2 | 11.394000 | 0.009079 | -0.224755 | 0.495351 | -1.753249 | 0.987674 | -0.161508 |
| DG | SC3 | 10.901000 | 0.022076 | 0.179322 | 0.732532 | -1.955549 | 0.983399 | -0.147636 |
| DG | SC4 | 6.459000 | 0.020184 | 4.177054 | 0.985317 | -5.043549 | 2.561237 | -0.392493 |
| DG | TE3 | 2.874000 | 0.001820 | 12.415070 | -7.473848 | -2.118647 | 2.501126 | -0.486522 |
| DG | TE5 | 8.036000 | 0.006764 | 4.659892 | 0.784825 | -6.864606 | 4.116754 | -0.722491 |
| DT | BB1 | 32.885000 | 0.082201 | -7.330068 | 2.166365 | -3.534657 | 2.314476 | -0.394454 |
| DT | BB2 | 3.806000 | -0.107230 | 9.566750 | -6.202361 | -0.495504 | 1.143006 | -0.214200 |
| DT | BB3 | -1.356000 | 0.567379 | 6.765954 | 4.089761 | -9.615125 | 4.409751 | -0.642398 |
| DT | SC1 | 5.951000 | -0.029265 | 2.596303 | -0.561522 | -1.565326 | 0.893228 | -0.141429 |
| DT | SC2 | 10.901000 | 0.021834 | 0.194630 | 0.723930 | -1.931995 | 0.968563 | -0.145126 |
| DT | SC3 | 4.314000 | -0.077456 | 12.498203 | -7.649942 | -3.003596 | 3.262633 | -0.644986 |
| DT | TE3 | 2.874000 | -0.002512 | 12.435764 | -7.553438 | -2.073635 | 2.512793 | -0.494371 |
| DT | TE5 | 8.036000 | 0.001199 | 4.917623 | 0.656370 | -7.233925 | 4.446366 | -0.794678 |
| A | BB1 | 32.885000 | 0.083391 | -7.360403 | 2.192064 | -3.565057 | 2.333236 | -0.397867 |
| A | BB2 | 3.806000 | -0.107298 | 9.589166 | -6.238736 | -0.482161 | 1.141293 | -0.213909 |
| A | BB3 | 3.594000 | 0.045373 | 9.591789 | -1.292022 | -7.108510 | 4.055712 | -0.633725 |

| | | | | | | | | |
|---|---|---|---|---|---|---|---|---|
| A | SC1 | 6.671000 | -0.008553 | 1.632224 | -0.064662 | -1.486942 | 0.785446 | -0.120835 |
| A | SC2 | 5.951000 | -0.026066 | 2.543999 | -0.484369 | -1.553574 | 0.864669 | -0.135090 |
| A | SC3 | 11.394000 | 0.008713 | -0.238913 | 0.489194 | -1.752894 | 0.992675 | -0.162913 |
| A | SC4 | 6.459000 | 0.019905 | 4.179750 | 0.976328 | -5.033298 | 2.555999 | -0.391550 |
| A | TE3 | 7.824000 | -0.048810 | 8.215579 | -0.894914 | -9.542937 | 6.331222 | -1.166729 |
| A | TE5 | 8.036000 | 0.016412 | 5.149022 | 0.834197 | -7.590683 | 4.520632 | -0.782608 |
| C | BB1 | 32.885000 | 0.083111 | -7.354321 | 2.186100 | -3.557883 | 2.329187 | -0.397200 |
| C | BB2 | 3.806000 | -0.108108 | 9.616792 | -6.287320 | -0.451266 | 1.133316 | -0.213253 |
| C | BB3 | 3.594000 | 0.044842 | 9.619198 | -1.335828 | -7.072004 | 4.039529 | -0.630982 |
| C | SC1 | 5.951000 | -0.029113 | 2.597004 | -0.555077 | -1.563446 | 0.889562 | -0.140613 |
| C | SC2 | 11.621000 | 0.013661 | -0.259592 | 0.489183 | -1.525505 | 0.836441 | -0.134073 |
| C | SC3 | 5.019000 | -0.032761 | 5.537769 | -0.951050 | -3.711308 | 2.161460 | -0.349186 |
| C | TE3 | 7.824000 | -0.058483 | 8.293199 | -1.125638 | -9.421976 | 6.354417 | -1.183569 |
| C | TE5 | 8.036000 | 0.004935 | 4.926220 | 0.648107 | -7.051000 | 4.260644 | -0.748191 |
| G | BB1 | 32.885000 | 0.083254 | -7.357360 | 2.189148 | -3.561548 | 2.331206 | -0.397523 |
| G | BB2 | 3.806000 | -0.107883 | 9.609308 | -6.274025 | -0.461927 | 1.137370 | -0.213831 |
| G | BB3 | 3.594000 | 0.045145 | 9.612347 | -1.315421 | -7.091505 | 4.047062 | -0.632010 |
| G | SC1 | 6.671000 | -0.008632 | 1.632523 | -0.065672 | -1.486805 | 0.785656 | -0.120889 |
| G | SC2 | 11.394000 | 0.009122 | -0.228690 | 0.496164 | -1.750390 | 0.986492 | -0.161416 |
| G | SC3 | 10.901000 | 0.022087 | 0.170328 | 0.732808 | -1.952920 | 0.983576 | -0.147909 |
| G | SC4 | 6.459000 | 0.020234 | 4.176650 | 0.987378 | -5.044199 | 2.561080 | -0.392438 |
| G | TE3 | 7.824000 | -0.051774 | 8.346067 | -1.029363 | -9.552119 | 6.377766 | -1.178980 |
| G | TE5 | 8.036000 | 0.005251 | 4.710706 | 0.667469 | -6.725387 | 4.036441 | -0.706057 |
| U | BB1 | 32.885000 | 0.083159 | -7.355311 | 2.187153 | -3.559038 | 2.330030 | -0.397385 |
| U | BB2 | 3.806000 | -0.107731 | 9.600999 | -6.261319 | -0.466683 | 1.136981 | -0.213516 |
| U | BB3 | 3.594000 | 0.045443 | 9.596259 | -1.292222 | -7.111432 | 4.056877 | -0.633828 |
| U | SC1 | 5.951000 | -0.029245 | 2.596687 | -0.561187 | -1.564771 | 0.892651 | -0.141308 |
| U | SC2 | 10.901000 | 0.021789 | 0.188390 | 0.722231 | -1.925816 | 0.966543 | -0.145013 |
| U | SC3 | 5.246000 | -0.045865 | 5.899781 | -1.506647 | -3.170544 | 1.937171 | -0.317010 |
| U | TE3 | 7.824000 | -0.029681 | 7.937832 | -0.330781 | -10.141202 | 6.633347 | -1.221112 |
| U | TE5 | 8.036000 | -0.009097 | 4.331935 | 0.434165 | -5.808314 | 3.524388 | -0.623824 |

**Table S4** Comparison of the protein/RNA models identified via metainference simulations, using atomistic (Kooshapur *et al.*, 2018) or Martini form factors. The agreement with SAXS data was measured with *CRYSOL* (Svergun *et al.*, 1995) using the maximum order of harmonics available and 18 points for the Fibonacci grid. The model quality was assessed using the Molprobity validation implemented in Phenix (Adams *et al.*, 2010; Davis *et al.*, 2007).

| | | **Refined Models** | |
|---|---|---|---|
| | | **Atomistic** | **Martini** |
| **Agreement with SAXS** | *CRYSOL* $\chi^2$ | 1.42 | 1.43 |
| | *CRYSOL* $\chi^2$ ($q$<0.3Å$^{-1}$) | 1.59 | 1.34 |
| **Model Quality** | Molprobity Score | 1.46 | 1.37 |
| | Clash-score | 0.58 | 1.15 |
| | Ramachandran favoured | 92% | 92% |
| | Ramachandran outliers | 2% | 2% |

**Figure S1** Calculated Martini form factors for DNA and RNA nucleotides. The grey lines represent the form factors back-calculated from each nucleotide bead in the library, while the coloured lines are the averages over all the individual form factors.

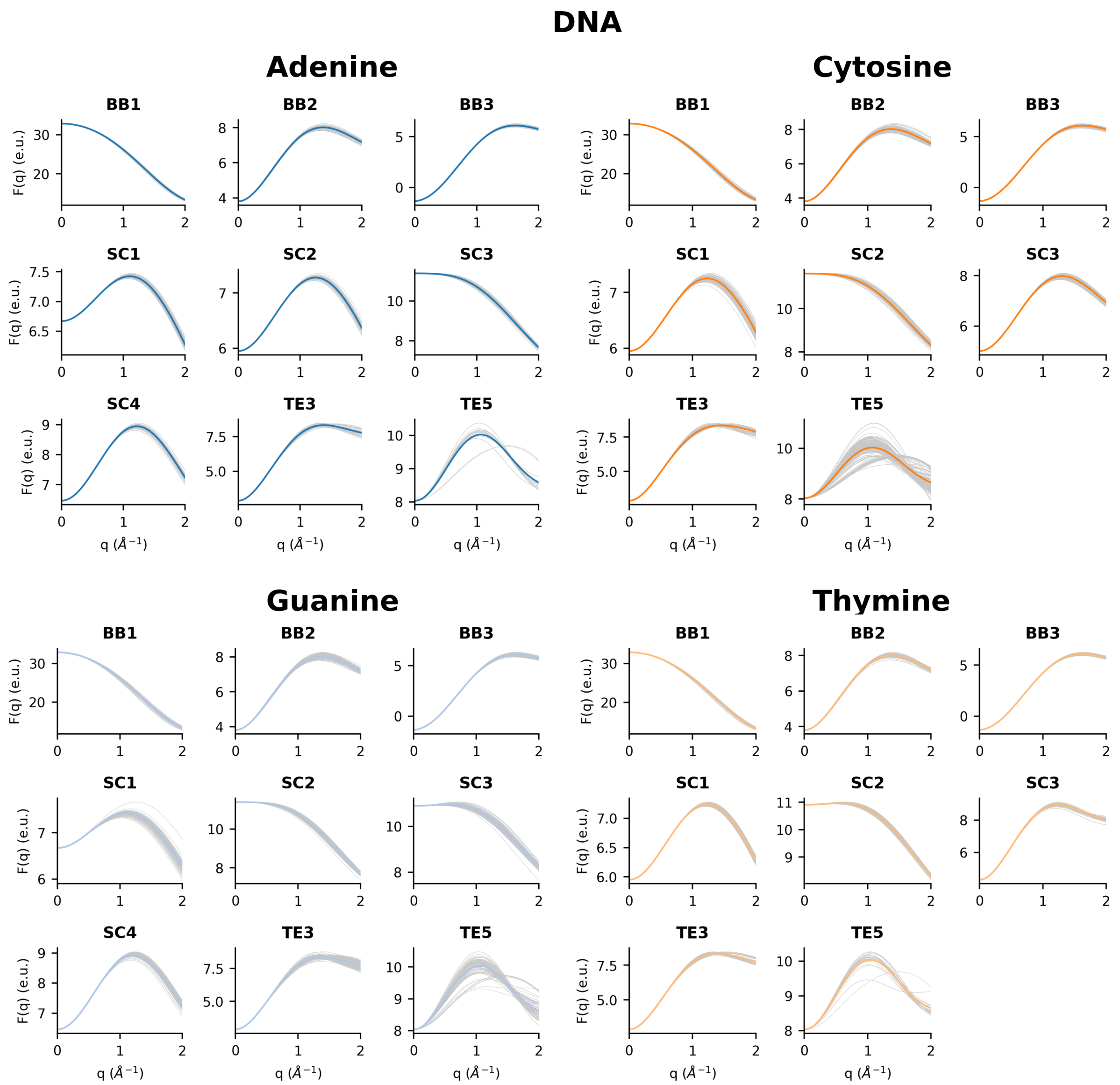

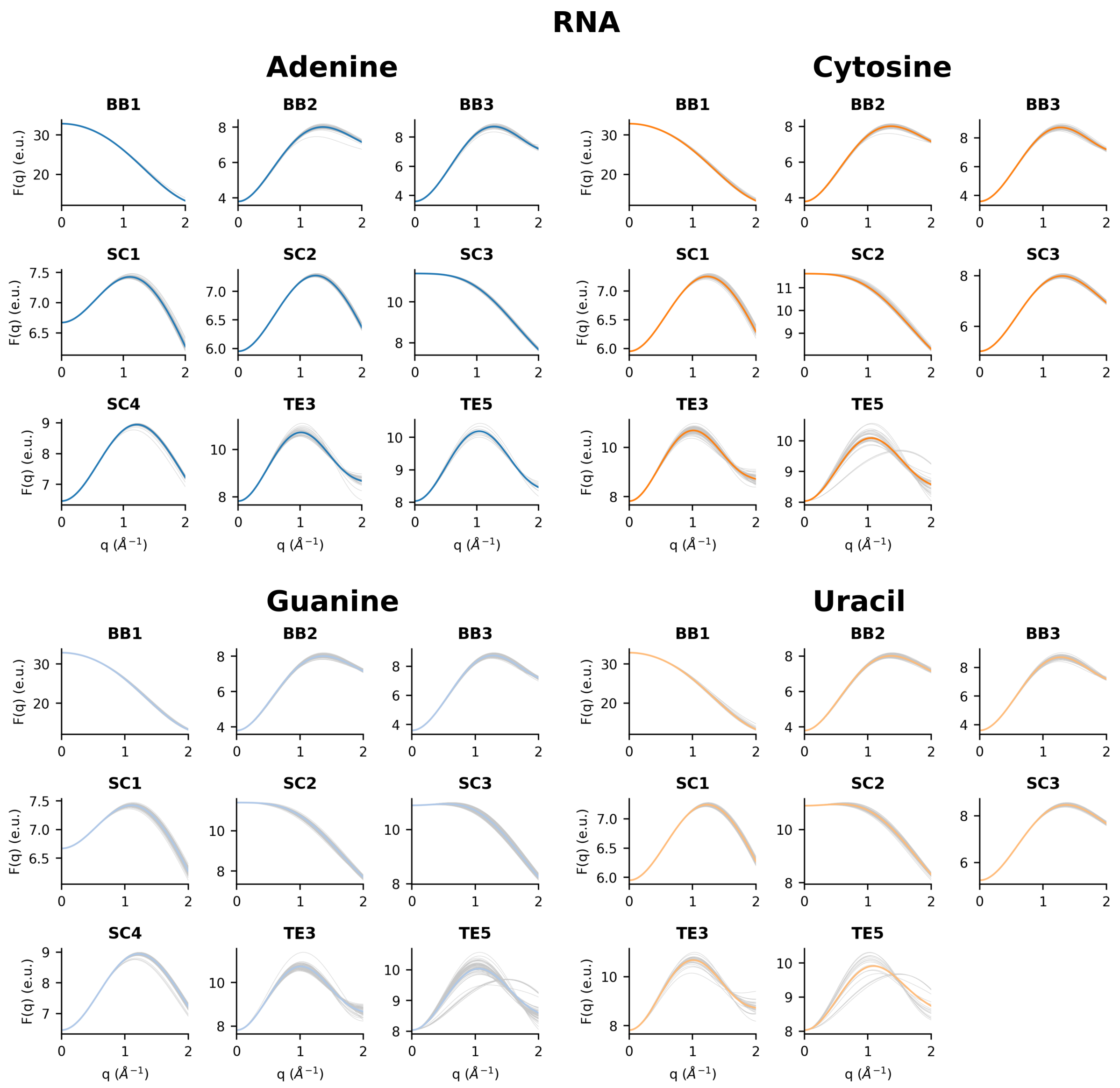
RNA
Adenine
Cytosine
Guanine
Uracil
BB1
BB2
BB3
SC1
SC2
SC3
SC4
TE3
TE5
F(q) (e.u.)
q (Å⁻¹)

**Figure S2** Distribution of $q_{threshold}$ values for 121 DNA crystallographic structures, coloured according to DNA classification. Average $q_{threshold}$ values are 0.56, 1.29, 0.51 and 0.78 Å$^{-1}$ for A-form, B-form, Z-form and quadruplex (Q) DNA, respectively.

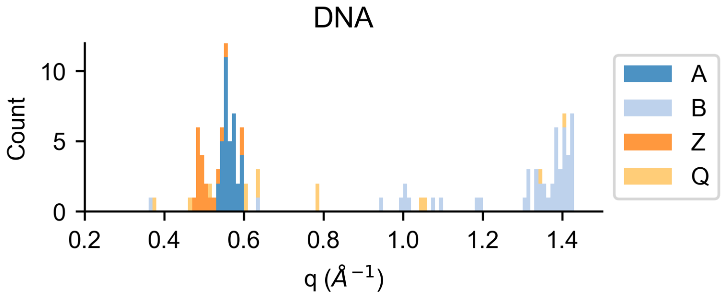


**Figure S3** For 121 DNA crystal structures are reported: (a) the distribution of *R*-values computed over a range of scattering vector *q* below 0.5 Å$^{-1}$, coloured according to DNA classification; (b) *R*-values, averaged over the DNA structures of X-form (with X indicating A-, B-, Z- form or quadruplex) and evaluated over a *q* range below a cut-off, as a function of the cut-off used.

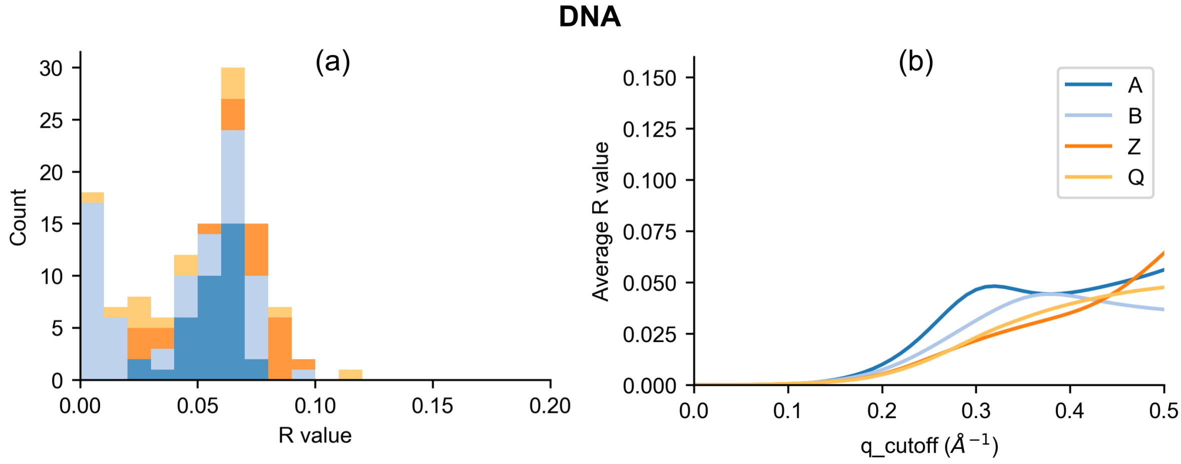


**Figure S4** Distribution of *R*-values computed over a range of scattering vector *q* below 0.45 Å$^{-1}$ for RNA (a) and DNA (b).

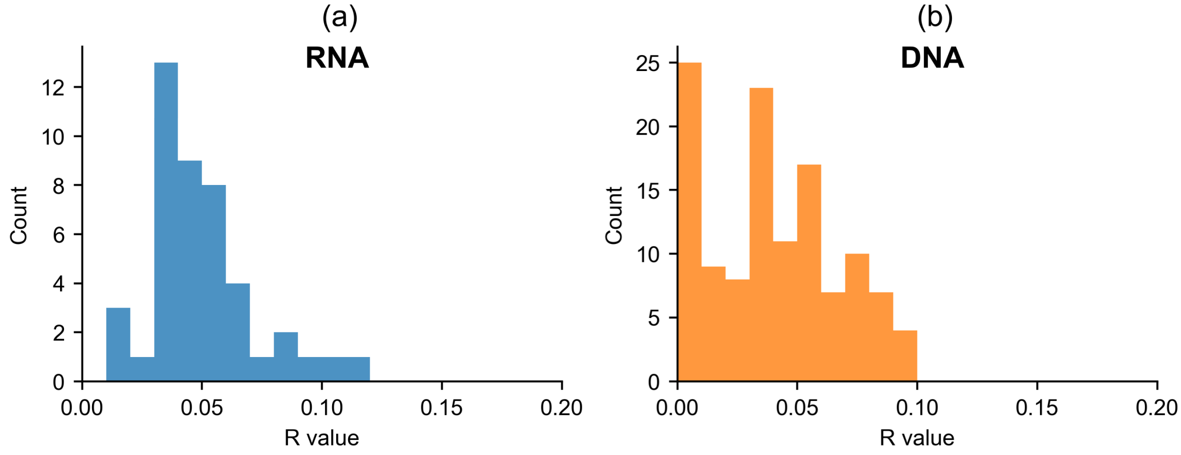
